# A novel human iPSC model of COL4A1/A2 small vessel disease unveils a key pathogenic role of matrix metalloproteinases in extracellular matrix abnormalities

**DOI:** 10.1101/2023.02.23.529680

**Authors:** Maha Al-Thani, Mary Goodwin-Trotman, Steven Bell, Krushangi Patel, Lauren K Fleming, Catheline Vilain, Marc Abramowicz, Stuart M Allan, Tao Wang, Zameel Cader, Karen Horsburgh, Tom Van Agtmael, Sanjay Sinha, Hugh S Markus, Alessandra Granata

**Author notes:** co-first author (Maha Al-Thani, Mary Goodwin-Trotman).

## Abstract

Cerebral small vessel disease (SVD) affects the small vessels in the brain and is a leading cause of stroke and dementia. Emerging evidence supports a role of the extracellular matrix (ECM), at the interface between blood and brain, in the progression of SVD pathology but this remains poorly characterized.

To address ECM role in SVD, we developed a co-culture model of mural and endothelial cells using human induced pluripotent stem cells from patients with *COL4A1/A2* SVD-related mutations. This model revealed that these mutations induce apoptosis, migration defects, ECM remodelling and transcriptome changes in mural cells. Importantly, these mural cell defects exert a detrimental effect on endothelial cells tight junctions through paracrine actions. *COL4A1/A2* models also express high levels of matrix metalloproteinases (MMP) and inhibiting MMP activity partially rescues the ECM abnormalities and mural cell phenotypic changes. These data provide a basis for targeting MMP as a therapeutic opportunity in SVD.

**Highlights:**

- A novel human iPSC-derived model of genetic SVD due to collagen IV (*COL4A1/A2*) mutations is described
- Mural cells expressing *COL4A1/A2* mutations have prominent ECM abnormalities as seen in patients and mouse models and contribute to endothelial cells defects
- ECM and endothelial cells abnormalities can be rescued by MMP inhibition in the *COL4A1/A2* model

## Introduction

Cerebral small vessel disease (SVD) is a leading cause of age-related cognitive decline and contributes to up to 45% of dementia cases worldwide (Gorelick et al., 2011). SVD is also responsible for 20% of ischemic strokes and is a common pathology underlying intracerebral haemorrhages (ICH) (Wardlaw et al., 2019). SVD refers to the sum of all pathological processes that affect the small vessels of the brain, including small arteries, arterioles and capillaries. With an aging population, SVD has major and growing global socio-economic impact (Lam et al., 2022). However, despite its importance, therapeutic approaches for SVD remain limited due to the lack of mechanistic understanding and relevant models required for target identification and drug discovery (Smith and Markus, 2020).

SVD features on neuroimaging include white matter hyperintensities, enlarged perivascular spaces, lacunar infarcts, microbleeds, microinfarcts, ICH and cerebral atrophy (Wardlaw et al., 2013). These features are associated with advancing age and several vascular risk factors, including hypertension and diabetes, though vascular risk factors do not account for all cases, leaving a large proportion of SVD unexplained (Wardlaw et al., 2014). Genetic factors have also been reported to be important with the identification of monogenic forms of SVD (Mancuso et al., 2020) and common variants which increase the risk of sporadic SVD (Chung et al., 2021.; Rannikmäe et al., 2015; Traylor et al., 2021). Dominant mutations in collagen type IV, a major component of the microvascular extracellular matrix (ECM), cause SVD presenting with both ICH and ischaemic (Gould et al., 2005; Jeanne et al., 2012). *COL4A1* and *COL4A2* mutations cause highly penetrant multi-system disorders by disrupting the ECM homeostasis and leading to ICH and porencephaly in human and mouse models (van Agtmael et al., 2005; Joutel and Faraci, 2014; Murray et al., 2014) referred as COL4A1 syndrome or Gould Syndrome. Importantly, both monogenic and sporadic forms of *COL4A*-related SVD are likely to share similar pathological mechanisms since rare coding variants in *COL4A1/A2* also occur in sporadic forms of ICH while common *COL4A1/A2* non-coding variants have been identified as risk factor for sporadic lacunar stroke (Chung et al., 2019; Persyn et al., 2020; Traylor et al., 2021), sporadic ICH (Malik et al., 2018; Rannikmäe et al., 2015) and white matter hyperintensities (Persyn et al., 2020) in the general population. This suggests that insights gained from a model of monogenic *COL4A1/A2* are likely to be relevant to common human SVD.

Although the mechanisms leading to SVD are ill-defined there is an emerging focus on the role of the ECM. The ECM of cerebral blood vessels is a key component at the interface between the cerebral microcirculation and the brain, providing structural support to the blood brain barrier (BBB) as well as influencing cell signalling and behaviour (Joutel et al., 2016). Genetic studies have revealed that most monogenic forms of SVD are caused by mutations in genes either encoding ECM proteins, or in proteins regulating ECM function (Joutel et al., 2016). In addition to extensive genetic support for a key role of the ECM in SVD, our recent work has found that genes related to SVD, including *COL4A1* and *COL4A2*, are significantly enriched in the cerebrovascular ECM network in both mouse and human brain (Pokhilko et al., 2021). To date, the mechanisms by which these ECM defects cause disease remains poorly understood. No analysis to date has been performed on the impact of these mutations on human endothelial and vascular smooth muscle cells, while the effects on ECM, BBB integrity and function remain undefined. This underscores the clear need for new models relevant to human SVD, which can be used to improve mechanistic understanding including the impact of microvascular ECM perturbations in SVD and to offer a platform for early evaluation of potential therapies.

Human induced pluripotent stem cells (hiPSC)-based ‘disease in a dish’ models are increasingly used as complementary tools to corroborate findings from animal model and provide a clinically relevant platform for drug screening (Vadodaria et al., 2020). To provide insights into the pathological mechanisms underlying COL4A1/Gould syndrome, which are still largely unknown, we established a SVD patient-specific *COL4A* iPSC model from two individuals with two representative glycine substitutions, one in *COL4A1* (G755R) and the other in *COL4A2 (G702D*) gene (Murray et al., 2014; Shah et al., 2010). We differentiated the hiPSC into neural crest-derived smooth muscle cells (mural cells) and endothelial cells and undertook phenotypic and functional assays and transcriptomic analysis.

## Results

### 1. Establishment and characterisation of COL4A1/A2 hiPSC-derived mural cells and endothelial cells

Two human iPSC lines (hiPSC) were used in this study: a *COL4A1^G755R^* with a G>A substitution in exon 30 of *COL4A1* gene resulting in a change from a glycine to arginine at position 755 (G755R) from a symptomatic patient and a *COL4A2^G702D^* with a G>A replacement in exon 28 of *COL4A2* gene, resulting in a change from a glycine to aspartic acid at position 702 (G702D) from the asymptomatic father of a patient (**Table S1)** (Murray et al., 2014; Shah et al., 2010). To control for genetic background, we generated isogenic corrected lines, in which the mutant allele (A) in *COL4A1* and *COL4A2* hiPSCs were substituted with the wild-type (WT) allele (G) (**Table S1-2; Figure S1A**). As further controls, we used three wild-type (WTs) hiPSC lines from healthy individuals, already established in the lab (**Table S1**). hiPSC lines were characterised for pluripotency markers expressions by immunostaining, qPCR profiling and formation of the 3-germ layers (**Figure S1B-D**). hiPSC were successfully differentiated into smooth muscle cells of neural crest origin (mural cells; MC) as previously described (Cheung et al., 2012; Serrano et al., 2019) **(Figure S2A)** and characterised for specific markers expression at neural crest stage (**Figure S2B, C**) and at fully differentiated stage (MC) by immunohistochemistry and qPCR (**Figure 1A-B; Figure S2D**).

**Figure 1.**
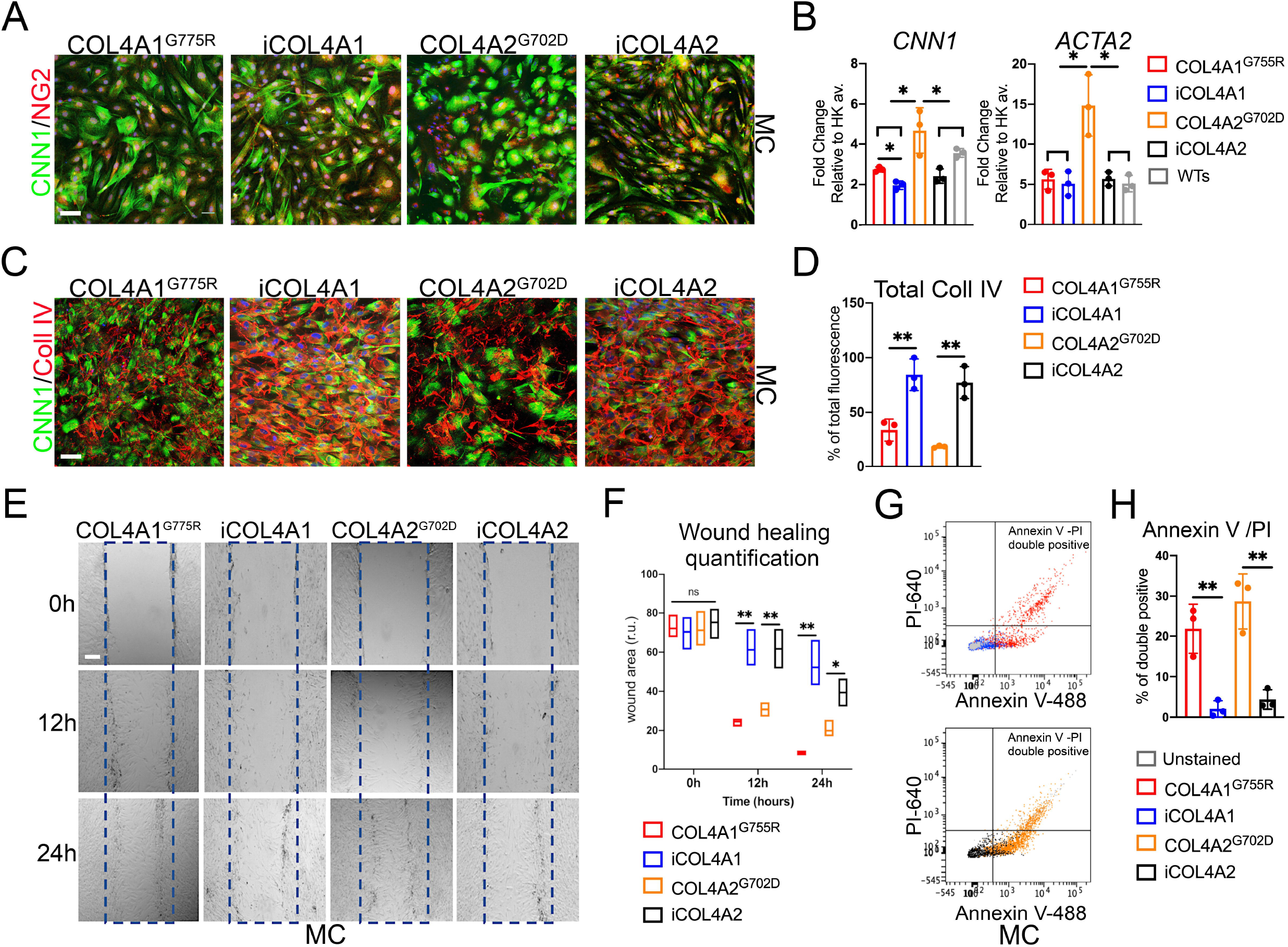
COL4A1^*G755R*^, COL4A2^*G702D*^ hiPSC-derived mural cells show abnormal collagen IV and phenotypic changes. **A)** Immunostaining for Calponin (CNN1) and Nerve/glial antigen 2 (NG2) in hiPSC-derived mural cells (MC) for COL4A1^*G755R*^, COL4A2 ^*G702D*^ and isogenic A1 (iCOL4A1) and A2 (iCOL4A2). **B)** RT-qPCR analysis showing increased expression for *CNN1* (COL4A1^*G755R*^ and COL4A2^*G702D*^) and *ACTA2* (COL4A2^*G702D*^) compared to controls: isogenics (iCOL4A1 and iCOL4A2) and WTs (three independent healthy control; **Table S1**). **C)** Immunostaining of extracellular collagen IV in non-permeabilized hiPSC-derived MC cultured in serumcontaining medium for 2 weeks, show significant decreased levels in COL4A1^*G755R*^ and COL4A2^*G702D*^ when quantified as % of total fluorescence **(D)** compared to isogenics. **E)** Representative images of scratch wound assays for hiPSC-MC and **(F)** quantification of the wound areas showing increased migration rate for COL4A1/A2 mutant MC compared to isogenic lines. **G)** Flow cytometric analysis of Annexin V-488 and propidium iodide (PI-640) in hiPSC-MC and **(H)** double staining quantification show higher apoptotic rate in COL4A1/A2 mutant compared to isogenic MC lines. Nuclei were stained with DAPI; scale bar=100μm. MC=neural crest-derived smooth muscle cells. The results are presented as means ± SD of three independent experiments; *\*P* < 0.05; \*\**P* < 0.01; ns, not significant. Statistical analysis was performed by 2-way ANOVA with Tukey’s multiple comparison test.

MC express both specific markers for smooth muscle cells (*CNN1, ACTA2, TAGLN*) and pericytes (*NG2* and *PDGFRA*), with the disease lines showing significantly increased expression levels for *CNN1* in *COL4A1/A2* and *ACTA2* in COL4A2^*G702D*^ (**Figure 1B**). MC are known to produce a variety of ECM proteins, including collagen IV. To assess collagen IV produced by the MC in the ECM, both *COL4A1/A2* disease and isogenic hiPSC-derived MC were stained with a specific antibody which recognised both collagen IV α1 and α2 chains (**Figure 1C**). There was a significant reduction in collagen IV staining in the disease *COL4A1/A2* mutant lines compared to the controls, as seen in patient fibroblasts (**Figure 1D)** (Murray et al., 2014). Moreover, hiPSC-derived MC with *COL4A1^G755R^* and *COL4A2^G702D^* have increased migration ability in the scratch assay (**Figure 1E-F).** They also exhibit higher apoptotic levels, when stained for Annexin V and Propidium iodide (PI) by flow cytometry compared to the isogenic controls (**Figure 1G-H)**, similar to previous findings from primary patient fibroblasts and skin biopsy (Murray et al., 2014). These data indicate that our MC recapitulate defects of *COL4A1/2* mutation and thus represent a valid model to explore disease mechanisms.

### 2. Mural cells contribute to the barrier phenotype in a co-culture and paracrine systems

Brain endothelial cells are known for their barrier function in the BBB, which may be compromised in SVD pathology (Hussain et al., 2021). However, the impact of collagen IV mutations on the BBB and cross talk between endothelial cells and MC remains very poorly understood. To assess this, *COL4A1/A2* disease and isogenic lines were differentiated into brain microvascular endothelial-like cells (BMEC) using a previously established protocol (**Figure S3A;** (Hollmann et al., 2017). These BMEC were characterised for their differentiation efficiency and expression of specific markers by qPCR and flow cytometry (**Figure S3B-D).** BMEC were then plated onto a 2D transwell setting alone or in presence of MC plated on the basolateral side and maintained for 6 days (**Figure 2A)**. During this time, daily readings were taken of transendothelial electrical resistance (TEER), a robust indicator of endothelial cell barrier integrity. The average of TEER measurement taken over the course of 6 days was expressed as a percentage relative to the peak value of the control for BMEC alone and in co-culture with MC were compared (**Figure 2B).** Isogenic control MC appear to promote barrier function by significantly increasing TEER values, while disease MC have little effect on *COL4A1/A2* BMEC TEER (**Figure 2B).**

**Figure 2.**
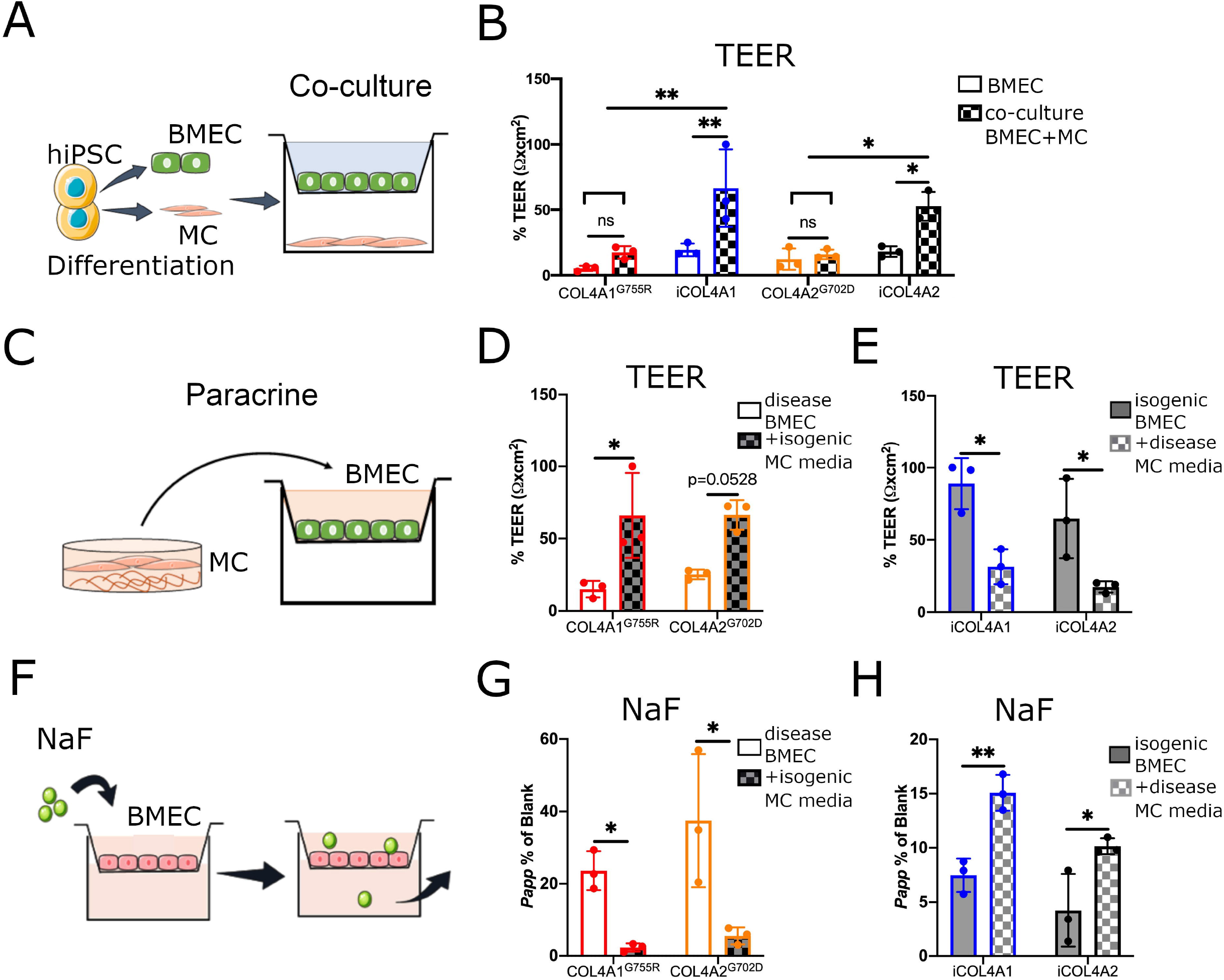
hiPSC-derived mural cells contribute to barrier function in a Transwell coculture system and COL4A1/A2 lines exert a detrimental effect. **A)** Schematic of co-culture with hiPSC-derived mural cells (MC) and brain microvascular endothelial-like cells (BMEC) in a Transwell device. **B)** Percentage of average transendothelial electrical resistance (TEER) measured across the 6 days relative to peak value of control expressed as resistance (Ω) x cm^2^ for hiPSC-derived BMEC in co-culture with MC increases compared to BMEC alone for isogenic COL4A1/A2. **C)** Schematic of the MC paracrine experiment with COL4A1^*G755R*^ and COL4A2^*G702D*^ BMEC TEER values benefiting from treatment with isogenic MC conditioned media **(D)**; while isogenic BMEC show decreased TEER values upon treatment with disease COL4A1/A2 MC conditioned media **(E). F)** Schematic of the sodium fluorescein (NaF) permeability assay in transwell setting. Isogenic MC paracrine effect positively reduce BMEC permeability in COL4A1^*G755R*^ and COL4A2^*G702D*^ lines after 6 days treatment **(G)**; while disease MC conditioned media treated isogenic BMEC show increased permeability to NaF compared to untreated BMEC **(H)**. TEER= transendothelial electrical resistance; NaF= sodium fluorescein; *Papp*=apparent permeability. The results are presented as means ± SD of three independent experiments; \**P* < 0.05; \*\**P* < 0.01; ns, not significant. Statistical analysis was performed by 2-way ANOVA with Tukey’s multiple comparison test.

Since in this setting, there is no cell-cell interaction between MC and BMEC, we further assessed the differential MC paracrine effect on the barrier properties, by treating isogenic BMEC with conditioned media of *COL4A1/A2* MC and vice versa for 6 days (**Figure 2C).** Interestingly, TEER values of disease BMEC tend to benefit from isogenic MC paracrine effect (**Figure 2D).** Conversely disease MC exert a paracrine effect by significantly decreasing TEER values in isogenic BMEC (**Figure 2E).** This paracrine effect of MC on barrier phenotype was confirmed by sodium fluorescein (NaF) size exclusion paracellular permeability assay (**Figure 2F)**, with the isogenic MC decreasing NaF permeability, thus promoting barrier tightness (**Figure 2G**), while disease MC appear to increase barrier permeability in isogenic BMEC (**Figure 2H)**. Collectively, these data show the secretome of disease MC to have detrimental effects on barrier function in *COL4A1/A2* associated SVD.

### 3. COL4A1/A2 mural cells affect endothelial tight junction levels and distribution via a paracrine effect

The integrity of tight junctions is essential for the BBB properties of brain endothelial cells (Nitta et al., 2003; Pan et al., 2017). Thus, to assess if tight junctions are affected in *COL4A1/A2* hiPSC-derived BMEC, we performed immunostaining analysis for the tight junction proteins, occludin and claudin-5 (**Figure 3A**). We observed striking discontinuities in occludin staining (white arrow) and frayed junctions evident with claudin-5 staining (white arrowhead) in *COL4A1^G755R^* and *COL4A2^G702D^* hiPSC-BMEC. These abnormalities were significantly more frequently in the mutant lines compared to isogenic controls (**Figure 3B)**. Moreover, this was associated with reduced occludin and claudin-5 total protein levels (**Figure 3C)**.

**Figure 3.**
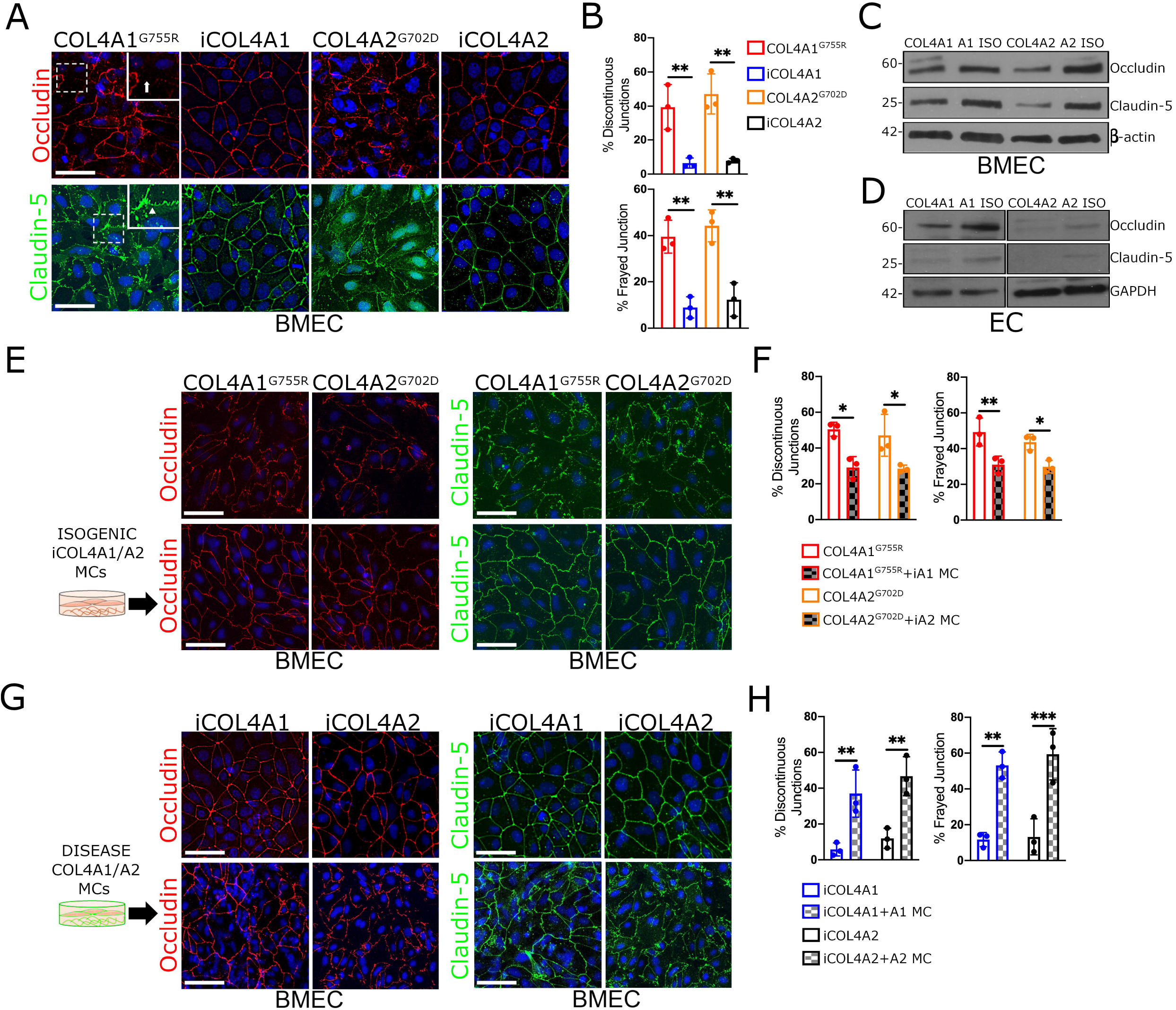
COL4A1/A2 mural cells contribute to endothelial tight junction abnormalities by paracrine effect. **A)** Junctional staining for occludin and claudin-5 in hiPSC-derived BMEC lines showing discontinuous junction (white arrow) and frayed junction (white arrowed) in zoom-in insert. **B)** Quantification of discontinuous and frayed junctions show higher percent in COL4A1^*G755R*^ and COL4A2^*G702D*^ lines compared to isogenic controls. Western blotting analysis show decreased total protein levels for occludin and claudin-5 in COL4A1^*G755R*^ and COL4A2^*G702D*^ derived BMEC **(C)** and iECs **(D)** compared to controls (A1 and A2 ISO). β-actin and GAPDH were used as loading controls. **E-F)** Immunostaining analysis of COL4A1^*G755R*^ and COL4A2^*G702D*^ BMEC tight junctions (occludin and claudin-5) upon 4 days treatment with isogenic MC conditioned media show less discontinuous and frayed junctions. **G-H)** Isogenic BMEC show increased percent of junctions abnormalities upon treatment with disease MC conditioned media. Nuclei were stained with DAPI; scale bar=100μm. The results are presented as means ± SD of three independent experiments; \**P* < 0.05; \*\**P* < 0.01; *** *P* < 0.001. Statistical analysis was performed by 2-way ANOVA with Tukey’s multiple comparison test.

To independently validate these findings and exclude they were due to the differentiation protocol, we adopted an alternative endothelial differentiation protocol to generate hiPSC-derived EC (iECS; **Figure S3E)** (Orlova et al., 2014a, 2014b). These iECs were characterised for expression of specific markers by flow cytometry and qPCR (**Figure S3E-G)** and were found to express lower levels of occludin and claudin-5 proteins in diseased lines compared to controls, validating our findings in hiPSC-BMEC (**Figure 3D**).

To assess if the levels and the distribution of occludin and claudin-5 in endothelial cells is regulated by the MC secretome, *COL4A1^G755R^* and *COL4A2^G 702D^* hiPSC-BMEC were treated with isogenic MC conditioned media for 4 days prior to immunostaining (**Figure 3E**). Notably, treatment with isogenic MC media significantly improved the presence of discontinuous and frayed junctions (**Figure 3F**). Conversely, a greater number of discontinuous and frayed junctions was observed when isogenic BMEC were treated with conditioned media from mutant *COL4A1/A2* MC (**Figure 3G-H).** These data clearly support that COL4 associated SVD includes tight junction defects in endothelial cells that are determined at least in part by a paracrine effect of the MC.

### 4. Transcriptomic analysis highlights ECM abnormalities in COL4A1/A2 mural cell lines

To identify potential mediators of the observed MC paracrine effects reported above, we performed a transcriptomic analysis on *COL4A1^G755R^* and *COL4A2^G702D^* and corresponding isogenic hiPSC-MC (**Figure 4A**). From the bulk RNA sequencing data, we identified 374 differentially expressed genes (DEGs) of which 283 were upregulated in the *COL4A* diseased lines. It also emerged that 59 DEGs were ECM proteins, and that matrix metalloproteinase (MMPs) were among the proteins misregulated (**Figure 4B; Table S5 and S6**). It is known that increased MMPs levels are associated with barrier disruption and stroke (Candelario-Jalil et al., 2011; Clark et al., 1997; Wallin et al., 2017). Thus, we proceeded to validate the transcriptomics findings by performing a MMP gene expression profiling using qRT-PCR. We observed increased *MMP2* and *MMP9* mRNA levels in both *COL4A1* and *A2* MC lines (**Figure 4C).** In addition, we also found a significant increase in *MMP14* mRNA levels in *COL4A1/A2* BMEC (**Figure 4D**). Interestingly, MMP14 which activates pro-MMP2, was also previously reported to be upregulated in aorta of mice with a *Col4a1* glycine mutation (*Col4a1*^+/SVC^ G1064D) that is a well-established model of Col4a1 associated SVD (**Figure 4D-E**; **Figure S4A)** (van Agtmael et al., 2005; Jones et al., 2016, 2019). The *MMP14* increase was also validated at protein levels in *COL4A1/A2* iECs (**Figure S4B).** These data clearly support that the ECM and MMPs are dysregulated in *COL4A1/A2* MC and vascular ECM.

**Figure 4.**
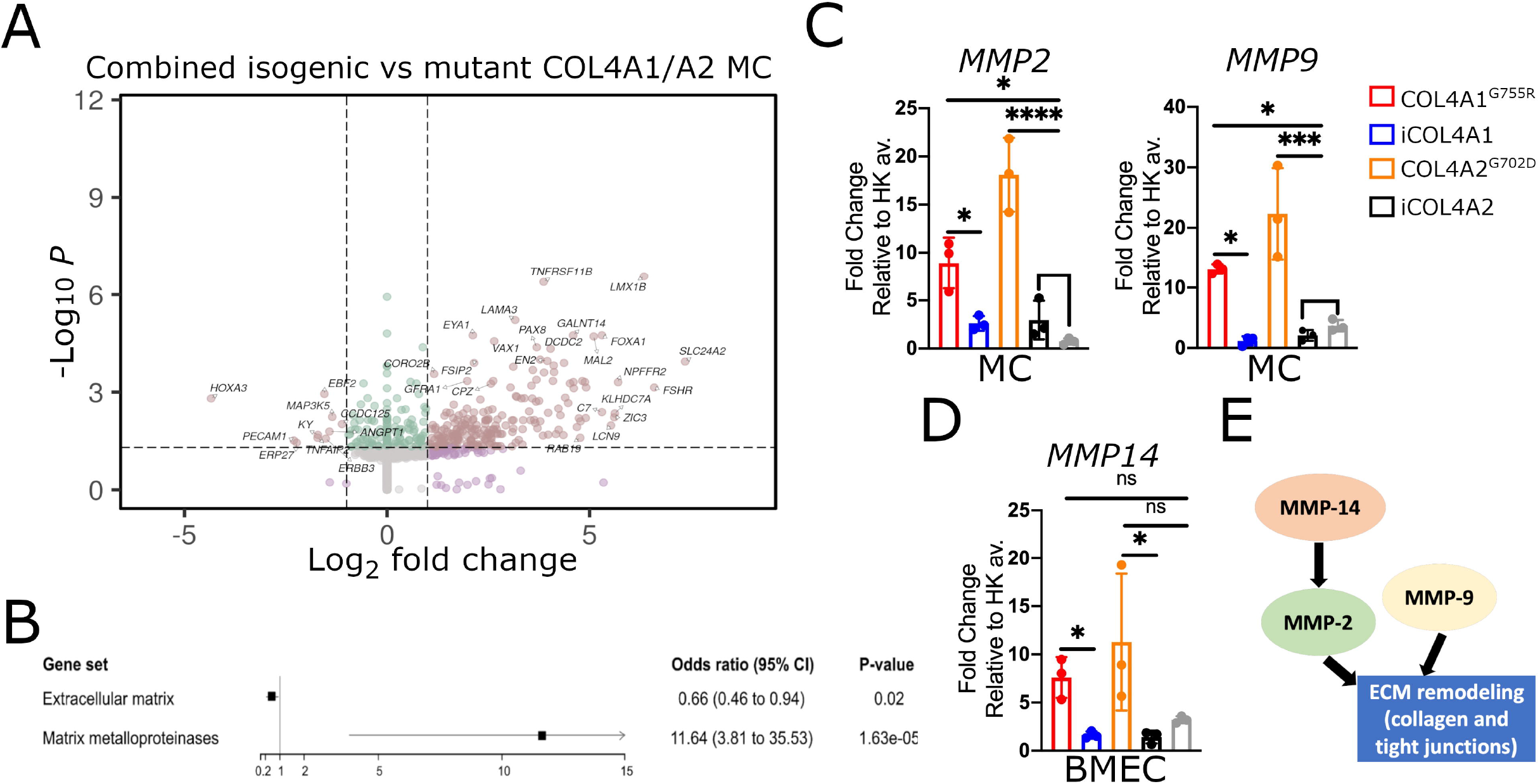
Transcriptomic analysis shows ECM abnormalities in COL4A1/A2 MC lines and MMPs upregulation. **A)** Volcano plot depicting differentially expressed genes in combined COL4A1^*G755R*^ and COL4A2^*G702D*^ compared to isogenic MC; matrisome protein are labelled and **B)** forest plot shows significant enrichment for ECM and MMPs in diseased MC**. C)** RT-qPCR analysis confirmed increased *MMP2* and *MMP9* mRNA levels in COL4A1^*G755R*^ and COL4A2^*G702D*^ MC. **D)** *MMP14* mRNA was found higher in COL4A1/A2 compared to isogenic hiPSC-BMEC lines and **E)** schematic of MMP2 −9 −14 contribution to ECM remodelling. The results are presented as means ± SD of three independent experiments; \**P* < 0.05; *** *P* < 0.001; **** *P* < 0.0001; ns, not significant. Statistical analysis was performed by 2-way ANOVA with Tukey’s multiple comparison test.

### 5. MMP inhibition rescues phenotypic alterations, ECM and tight junction defects

Since, MMPs are important for matrix remodelling, including collagens, and because they also target tight junctions for degradation, we hypothesized that MMPs could mediate the *COL4A1/A2* ECM phenotype seen in our *in vitro* model. Thus, we proceeded to treat *COL4A1/A2* hiPSC-derived BMEC with the pan-MMPs inhibitor, doxycycline (DOXY), which appears to successfully represses MMP2 and MMP9 activity after 72 hours treatment by zymography (**Figure 5A**). Upon 4 days treatment with 8μM DOXY, diseased BMEC stained for occludin and claudin-5 show a significant reduction of discontinuous and frayed junction percentage compared to control (DMSO) (**Figure 5B-C).** Similarly, DOXY mediated MMPs inhibition improved tight junction abnormalities in *COL4A1^G755R^* and *COL4A2^G702D^* hiPSC-ECs (**Figure S4C-D).** In addition, treatment with DOXY appeared to increase total occludin and claudin-5 protein levels by western blotting (**Figure 5D).** Remarkably, DOXY treatment benefited on the barrier properties of the BMEC as evidenced by significantly increasing TEER values (**Figure 5E**) and reducing the NaFl permeability percentage (**Figure 5F).**

**Figure 5.**
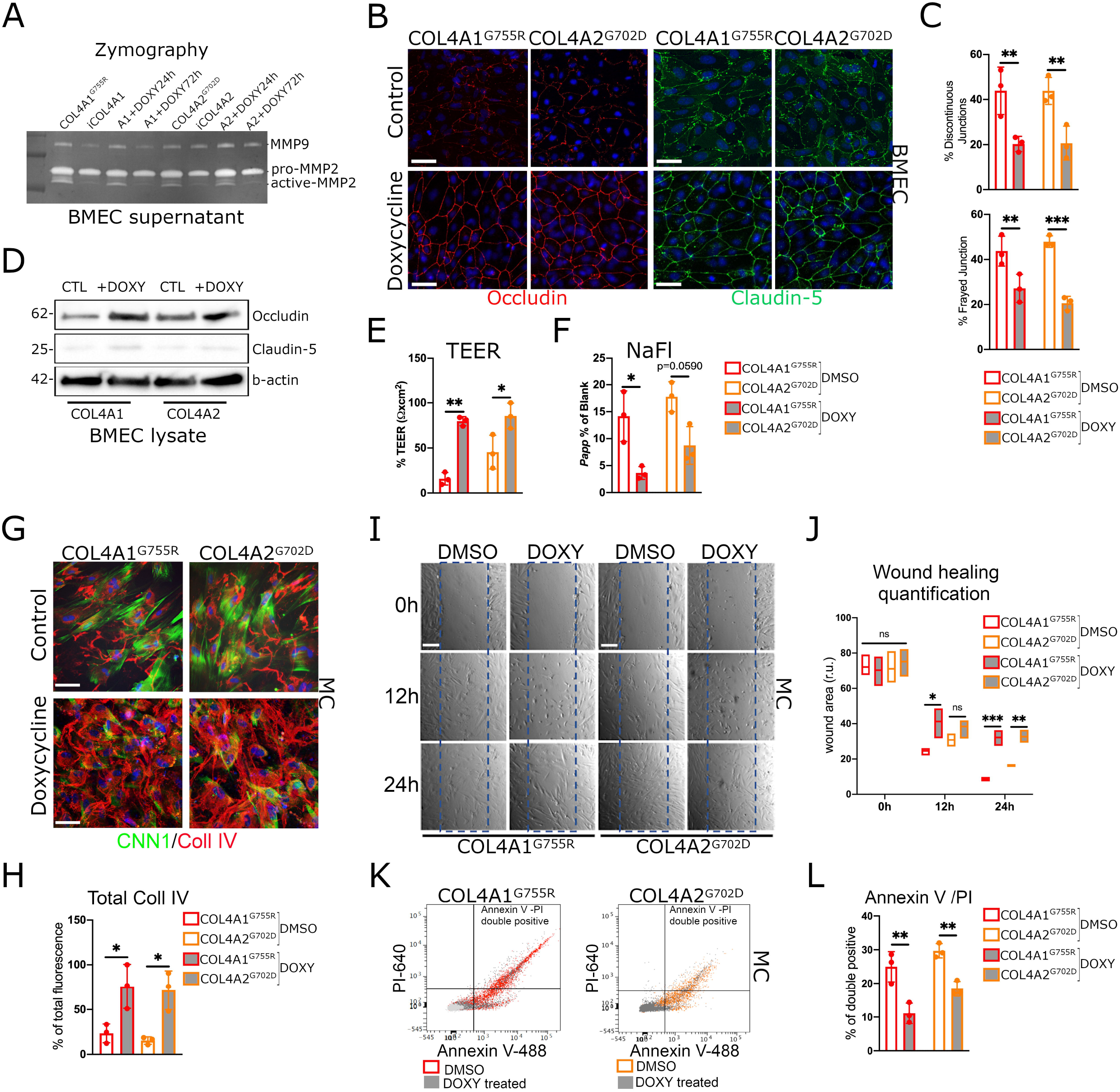
Doxycycline treatment ameliorates tight junction abnormalities and reverts COL4A1/A2 MC collagen IV defect and phenotypic changes. **A)** Zymography analysis of BMEC supernatants show higher MMP9 and MMP2 activity levels in mutant *COL4A1/A2* compared to isogenics, which decrease in response to doxycycline (8 uM) mediated MMPs inhibition. **B-C)** Immunostaining analysis quantification of occludin and claudin-5 in COL4A1^*G755R*^ and COL4A2^*G702D*^ BMEC treated with doxycycline (DOXY) for 4 days show lower percent of tight junction abnormalities (discontinuous and frayed junctions) compared to untreated control (DMSO). **D)** Protein blot analysis shows increased occludin and claudin-5 levels upon treatment with DOXY. β-actin was used as loading control. Doxycycline treatment improves both TEER **(E)** and NaF **(F)** readouts in *COL4A1/A2* mutant BMEC compared to untreated controls (DMSO). **G)** Immunostaining analysis of collagen IV in *COL4A1^G755R^* and *COL4A2^G702D^* MC treated for 4 days with doxycycline (DOXY; 10uM), and total fluorescence quantification **(H)** show higher percent in ECM compared to control (DMSO). **I)** Representative image of scratch assay for *COL4A1/A2* hiPSC-derived MC control (DMSO) and doxycycline treated (DOXY) and wound area quantification **(J)** show lower migration rate upon treatment with DOXY compared to controls. **K-L)** DOXY treatment improves apoptotic levels in *COL4A1^G755R^* and *COL4A2^G702D^* MC compared to untreated. The results are presented as means ± SD of three independent experiments; \**P* < 0.05; \*\**P* < 0.01; *** *P* < 0.001; ns, not significant.

We also looked at the effect of inhibiting MMPs by DOXY treatment (10uM) on collagen IV deposition in *COL4A1/A2* hiPSC-derived MC and we found that collagen IV total fluorescence levels detected by immunostaining in the *COL4A1/A2* ECM increased upon treatment compared to control (**Figure 5G-H**). DOXY treated diseased MC cells also show a significant decreased migration rate at 24 hours (**Figure 5I-J)**, and lower apoptotic levels (**Figure 5K-L)** comparable to isogenic controls (**Figure 1E-H).** These data establish a role for ECM remodelling due to MMPs caused by *COL4A1/A2* mutations and provide *in vitro* evidence that modulating MMPs may represent a therapeutic target for SVD.

## Discussion

There is a critical need to develop new relevant models relevant to human SVD to provide mechanistic insights as well as a foundation to test potential treatments for this debilitating disorder. To address this, we characterised a novel *in vitro* model of human SVD produced by differentiating iPSC generated from patients with a *COL4A1* or a *COL4A2* SVD related mutation into MC. Using this new model, we show phenotypic differences caused by *COL4A1/A2* mutations and identify a key role for ECM remodelling linked to BBB alterations. Importantly we show that these deleterious effects can be partially reversed through inhibition of MMPs.

We used *COL4A1/A2* patient derived hiPSC-MC in a co-culture system with brain microvascular endothelial-like cells to mimic the changes seen in patients’ small vessels and to investigate underlying pathological mechanisms. Firstly, we observed increased expression of smooth muscle cells markers, such as *CNN1* and *ACTA2*, in *COL4A1/A2* hiPSC-derived MC, which may suggest hypermuscularization as previously shown in a Col4a1 mouse model (Ratelade et al., 2020). Disease mural cells also showed an ECM defect including lower levels of extracellular collagen IV in agreement with previous findings from patient cells (Jeanne et al., 2015; Murray et al., 2014). We also determined phenotypic changes in disease mural cells, including increased migration and apoptotic rates which parallel previous studies using primary patient fibroblasts and skin biopsy samples (Murray et al., 2014). A loss of smooth muscle cells has been reported before and could be caused by several mechanisms, including ECM remodelling and endoplasmic reticulum stress (Ratelade et al., 2018, 2020).

Given the key strategic location of the ECM at the interface between blood and brain, a central aim of the study was to determine where COL4 disease influences barrier related properties. Interestingly, *COL4A1/A2* patient-derived MC exerted a detrimental effect on the endothelial barrier functions by a paracrine effect-evidenced by our transwell setup and with MC conditioned media treatment.

In this study, for the first time, we provided insight into the transcriptional features of *COL4A1/A2* patient-derived MC. Strikingly 15% of changes affected ECM proteins, including MMPs. Collagen IV is a substrate for the proteolytic activity of the gelatinases MMP2 and MMP9 and increased MMP2 and MMP9 expression has been associated with breakdown of collagen type IV in both human and animal models (Roach et al., 2002; Rosell et al., 2008), as well as with degradation and cellular rearrangement of the endothelial tight junctions (Bauer et al., 2010; Liu et al., 2012; Yang et al., 2007). Moreover, MMPs are known to play a role in smooth muscle migratory behaviour and may facilitate MC migration in our *COL4A1/A2* model by promoting proteolysis of the ECM proteins. For instance, it has been shown that MMP9 regulate pericyte migration in a rat model of cerebral ischemia, and downregulation of MMP9 after ischemia-reperfusion, appears to inhibit the migration of pericytes contributing to blood brain barrier integrity (Underly et al., 2017).

These findings suggest MMPs could play a role in the ECM alterations in *COL4A1/A2* related SVD and could present a novel therapeutic opportunity. In support of this, targeting MMPs using the pan-MMP inhibitor, doxycycline, partially rescued the disease MC phenotypes, including promoting collagen IV extracellular levels, reducing migration and apoptotic levels, and improving BMEC/iECs tight junction abnormalities. In other studies, doxycycline was shown to reduced vascular remodelling and damage induced by cerebral ischemia in a stroke animal model, the stroke-prone spontaneously hypertensive rats (Pires et al., 2011). However, doxycycline is a broad spectrum MMP inhibitor with potentially significant side effects. It would therefore be preferable to generate more selective small molecules and/or antibodies to inhibit specific MMP. Our model offers a human physiological platform for novel target discovery and to screen for more selective therapies in SVD.

Overall, major strengths of this work are 1) the generation of a new hiPSC-derived disease model for SVD, which replicates phenotypic changes observed in patients and *Col4a1* animal model, including ECM abnormalities; 2) this disease-relevant model can be used as new tool for the analysis of signalling pathways to identify therapeutic targets and 3) to screen and test for potential drugs against SVD.

Our work has limitations. Generating representative brain endothelial cells that possess endothelial identity while replicating the BBB properties, including elevated TEER and small molecule low permeability, has been a challenge highlighted in recent hiPSC work (Lu et al., 2021). We initially used the Hollman *et al* protocol, which originates from the Lippman lab (Hollmann et al., 2017). These cells display high TEER, however, they do also express epithelial-related genes and lack angiogenic properties (Lu et al., 2021). In view of these limitations, we then successfully validated our results using a generic endothelial protocol (Orlova et al., 2014b), however this lacks barrier-like functions. Further research is required to improve the current protocols for generation of BMEC cells, based on the emerging understanding of the BBB from single-cells sequencing studies (Garcia et al., 2022). An additional future direction of this work could involve the co-culturing of endothelial cells with pericytes and astrocytes in a 3D microfluidic model to promote endothelial barrier properties, and more closely replicate the *in vivo* microenvironment of the cerebrovasculature.

In conclusion, our novel hiPSC-derived mural cells model of *COL4A1/A2* mutations, supports a key role of the ECM in SVD and suggests that targeting ECM-related proteins like MMPs may be a promising potential therapeutic option.

## Experimental procedures

Tables and additional methodology are provided in the ***Supplemental Information*.**

### HiPSC culture

All the hiPSC lines use for this study are listed in **Table S1.** WT hiPSC lines were purchased from the HiPSci Human stem cell initiative cell bank (https://www.hipsci.org). *COL4A1^G755R^* hiPSC line was generated from skin biopsy from a SVD patient, recruited at the Stroke Research Group at the University of Cambridge and reprogrammed by the Cambridge iPSC core and two independent clones were used for this study. COL4A2^G702D^ hiPSC line was obtained by Professor Tom Van Agtmael (Murray et al., 2014). Isogenic control line for *COL4A1^G755R^* (iCOL4A1) and *COL4A2^G702D^* (iCOL4A2) were generated by CRISPR-gene editing method as described below and two independent clones were used for each line (**Figure S1A-B)**. *COL4A1/A2* mutant and isogenic hiPSC lines were characterised for pluripotency markers expression by immunostaining and qPCR (**Figure S1C-D)** and by formation of the 3 germ-layers (**Figure S1E).** All hiPSC lines were cultured in TeSR™-E8 media (STEMCELL Technologies) or E8 media (Dulbecco’s Modified Eagle Medium/Nutrient Mixture F-12 (DMEM/F-12) with Insulin-Transferrin-Selenium (Thermo Fisher Scientific), Sodium Bicarbonate (Thermo Fisher Scientific), and L-ascorbic acid (Merck) supplemented with FGF2 (4 ug/mL; Biochemistry Department, University of Cambridge) and TGF-β1 (1.74 ug/mL; R&D Systems) using Vitronectin XF (STEMCELL Technologies) as chemically defined xenofree cell culture matrix. All hiPSC lines were validated by the Cambridge Biomedical Research Centre iPSC core and routinely tested for presence of mycoplasma contamination by Mycoplasma Experience LTD.

### Transwell co-culture

Either 12-well or 24-well Transwells^®^ (Corning^®^ 0.4 μm pore; Sigma Aldrich) were coated on the apical and basolateral side with collagen IV/fibronectin. hiPSC-mural cells were dissociated with TrypLE and seeded onto the plate bottom of the Transwell^®^ coated with 0.1% Gelatin. After incubation for 1 hour, hiPSC-BMEC were dissociated and the seeded onto the apical side. The next day, Transwells^®^ with BMEC with(out) mural cells were maintained without any further medium changes for up to 6 days before analyses.

#### Paracrine

MC were serum starved for 4 days. At day 5, the MC serum-starved conditioned media was added to BMEC seeded onto collagen IV/fibronectin coated 24-well Transwells^®^ for TEER and NaFl analyses or 24-well plates for immunostaining assay. Initial TEER measurement was taken after 24h and afterwards on daily basis. For NaFl and immunostaining assays, BMEC were treated with condition media, refreshed every other day, for 6 days.

#### BMEC functional assays

##### TEER

Transendothelial Electrical Resistance (TEER) measurements were taken every 24 hours, from day 1 to day 6 of subculture of BMEC onto Transwells^®^ using an EVOM2 Voltohmmeter/STX2 electrodes (World Precision Instruments). The STX2 electrode was positioned within the well and the resistance (Ω) was recorded three times to calculate the mean resistance. All values are given as Ωxcm^2^ after subtracting the resistance of an empty coated Transwell^®^ well maintained in the same culture media (blank) and multiplying by the surface area (0.33cm^2^) as described previously (Lee et al., 2018). TEER was expressed as percentage of the average measurement across the 6 days relative to peak value of control.

##### NaFl

At 2 days post-subculture of BMEC onto 24 well Transwells^®^, spent media was removed from the upper chamber of the Transwell^®^ and replaced with 600μl of Sodium Fluorescein (NaFl, Sigma-Aldrich 1mg/ml) diluted 1:100 in endothelial serum-free media with B27. Samples of 100μl were taken from the basolateral side every two hours for eight hours. Raw fluorescence was measured with a TECAN Infinite M200 Pro plate reader (excitation wavelength of 460nm and emission 515nm; gain of 50, 25 flashes; z-position of 20000). Quantification was represented as percentage of total fluorescence relative to empty coated Transwell^®^ well (blank) as previously (Lee et al., 2018). The following formulae were used:

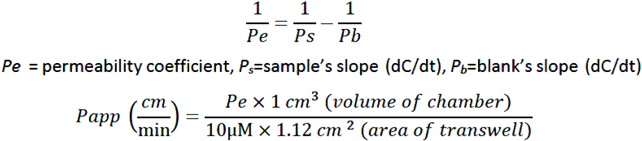

#### Doxycycline treatment

hiPSC-derived Mural cells were treated with doxycycline (10uM; Sigma) in DMEM+10% FBS for 4 days, with media change every other day, and then harvested for analyses. BMEC/iECs were treated with 8uM of doxycycline in EC medium, with media change every other day, and collected at 24h and 72h for zymography and at day 6 for immunostaining and western blotting analysis. TEER measurements were taken every 24 hours from day 1 to day 6 of subculture of BMEC onto Transwells in media supplemented with doxycycline. NaF permeability assay was performed after 6 days of doxycycline treatment.

#### Zymography

Gelatin zymography was performed in 10% acrylamide gels containing gelatin (4mg/ml; Novex 10% Zymogram gels; ThermoFisher). SDS-PAGE was performed using Tris-glycin**e** SDS sample and running buffers as described by the manufacturer. After electrophoresis, SDS was replaced by Triton X-100 (2.5%), thus renaturing gelatinases. Gels were then incubated in Tris buffer containing NaCl and ZnCl_2_ at 37°C for 24 h. Gels were then stained with Coomassie Blue solution, followed by de-staining (Methanol, Acetic Acid solution) and visualised by using Gel Doc™ XR+ system (BioRad).

##### Annexin V apoptosis assay

1 × 10^6^ cells/ml were harvested and resuspended in 1× annexin-binding buffer and incubated with 5 μl of Annexin V–488 (Alexa Fluor 488 Annexin V/Dead Cell Apoptosis Kit; Life technologies) for 15 min at room temperature. Cells were then resuspended in PBS with the addition of propidium iodide (PI 1:300) and measured with a BD LRSFortessa Flow cytometer. Flow cytometric data were analysed with FCSalyzer 0.9.15-alpha software.

#### RNA sequencing

##### Samples preparation

Three sets (biological replicates) of hiPSC-derived mural cells grown in DMEM+10%serum for 2 weeks were harvested. Total RNA was isolated from cells using RNeasy Mini Kit (QIAGEN) following the manufacturer’s instructions. Purified RNA samples were quantified and qualified using Nanodrop One (Thermo Scientific). Upon ribosomal RNA depletion, libraries were prepared using a NEBNext RNA library Prep kit (Illumina). The samples were run on an Novaseq6000 S4 lane and150 bp paired-end reads were generated.

##### Data analysis

The resulting base call files were converted to fastq files using the bcl2fastq program with default parameters. Alignment in STAR (2.7.10a) using a modified version of the ENCODE-DCC RNAseq pipeline annotated using GENCODE v39 (hg38) was performed (Dobin et al., 2013). An overhang of ±150bp around the annotated junction was used to construct the splice junction database. Gene-level RNA expression quantification was performed with RSEM (Li and Dewey, 2011).

Differential expression analyses were carried out using DESeq2 in R v4.0.4 (Love et al., 2014). We specified a false discovery rate of 5% and applied a Bayesian shrinkage estimator to effect sizes using approximation of the posterior for individual coefficients. Results were visualised using the EnhancedVolcano (1.12.0) package.

Enrichment of gene-sets of interest was calculated using logistic regression (i.e., whether a gene was statistically significantly differentially expressed). We used data from human samples to categorise genes associated with the extracellular matrix (ECM) (Pokhilko et al., 2021) and matrix metalloproteinases (**Table S5**). Pathways enrichment analysis was performed using the Reactome (**Table S6**) (Wu and Haw, 2017). We chose a 5% FDR to indicate statistical significance.

#### Statistical analysis

Statistical analysis was calculated using GraphPad Prism 9.00 (GraphPad Software Inc.). Two-way analysis of variance (Tukey’s multiple comparison test) was used to determine statistically significant differences between the groups. Results are presented as standard deviation (SD). All experiments represent the results of at least three independent biological replicates (measurements of biologically distinct samples). *P < 0.05; **P < 0.01; ***P < 0.001.

## Author contribution

AG contribute to the conception, design and interpretation of the hiPSC model data and to the drafting of the article. MA and MGT equally contributed to the acquisition and analysis of the data. SB undertook the transcriptomic data analysis. KP helped with hiPSC differentiation and establishing the co-culture and paracrine model. TVA and LKF undertook the mouse aorta dissection and analysis, KH and TVA provided critical reading of the manuscript. CV and MA are the clinicians for the patient with the *COL4A2* mutation. HSM is the clinician for the patient with *COL4A1* mutation, contributed to the revision of the article and the supervision of all studies. All authors contributed to the article and approved the submitted version.

## Acknowledgments

We thank the NIHR Cambridge BRC Cell Phenotyping Hub for the assistant with the Leica SP5 microscope and the Flow cytometry core facilities at the Cambridge Institute for Medical Research (CIMR). We also thank Professor L. Vallier and the hiPSC core facility for generating the *COL4A1* hiPSC line.

