## Supplemental Information for "A novel human iPSC model of COL4A1/A2 small vessel disease unveils a key pathogenic role of matrix metalloproteinases in extracellular matrix abnormalities"

***Supplemental Experimental procedures***

*HiPSC differentiation into NC-SMC*

For NC differentiation, hiPSC were detached from Vitronectin coated plates using ReLeSR (STEMCELL Technologies) as previously described (Cheung et al., 2012; Serrano et al., 2019). Clumps were plated at a density of 300 in 0.1% gelatin-coated six well plates in CDM-polyvinyl alcohol (PVA) for 4 days without splitting. CDM was composed of Iscove's modified Dulbecco's medium plus Ham's F12 NUT-MIX (Thermo Fisher Scientific) medium in a 1:1 ratio, supplemented with chemically defined lipid concentrate (Thermo Fisher Scientific), transferrin (Roche Diagnostics), insulin (Roche Diagnostics), and monothioglycerol (Sigma) supplemented with FGF2 (12 ng/mL; R&D Systems) and SB-431542 (10 mmol/L; Tocris), referred as FSB. After 4 days, hiPSC was dissociated using TrypLE Express (Gibco) and seeded as single cells at a 1:3 ratio on 0.1% gelatin-coated plates in FSB. NC cells were passaged every time reached confluence, up to 12 passages.

For NC-SMC differentiation, NC cells were dissociated using TrypLE Express and cultured in SMC differentiation medium (CDM-PVA supplemented with PDGF-BB (10 ng/ml, Peprotech) and TGF-β1 (2ng/ml, Peprotech) for 12 d. For long-term cultures, SMCs were subsequently grown in MEM (Sigma-Aldrich M5650) containing 10% FBS (Sigma-Aldrich F7524) up to 4 weeks.

*HiPSC differentiation into BMEC and iECs*

hiPSCs were differentiated to brain microvascular endothelial-like cells (BMEC) as previously described, with minor modifications (Hollmann et al., 2017). hiPSCs were washed once with 1X PBS (Corning®), dissociated with StemPro™ Accutase™ Cell Dissociation Reagent (Thermo Fisher Scientific) for 4 minutes, and collected by centrifugation. hiPSCs were then resuspended in E8 medium containing 10 μM Y27632 (Tocris Bioscience) and seeded onto Matrigel-coated 6-well plates at a density of $1.56x10^4$ /$cm^2$. The following day, the cells were switched to TeSR™-E6 medium (Stem Cell Technologies) (Lippmann et al., 2014) to initiate differentiation. Media was changed every day for 4 days. On day 5, the cells were switched to Endothelial media (EC), which consisted of a basal human endothelial serum-free media (SFM; Thermo Fisher Scientific), supplemented with B27 (Fisher Scientific), basic fibroblast growth factor (bFGF; 10ng/ml; R&D Systems) and all-trans retinoic acid (RA; 10μM; Sigma-Aldrich). Cells were then left to incubate for 48 hours in EC medium without a media exchange. On day 6, resultant BMEC cells were washed with PBS and dissociated with accutase to single cells and plated at a density of $5.0x10^6$ /well on 12 well culture plate or 24-well Transwells (CLS3470, Corning®) coated with collagen IV (from human placenta, 1mg/ml, Bornstein, and Traub Type IV; C5533, Sigma Aldrich) and fibronectin (from bovine plasma, 1mg/ml, F1141, Sigma Aldrich). 24 hours after plating, media was refreshed to EC medium without bFGF and RA. Subsequent media changes were performed every 2 days for 6 days.

hiPSC-ECs (iECs) were differentiated using a previously reported protocol with minor modifications (Orlova et al., 2014b). Briefly, hiPSCs were maintained in TeSR™-E8 medium on vitronectin-coated 6-well plates and seeded at day-1. Twenty-four hours after seeding E8 medium was replaced with B(P)EL medium supplemented with 8 μM CHIR. On day 3, the medium was replaced with B(P)EL medium supplemented with VEGF-A (50 ng/ml; Peprotech) and SB431542 (10 μM; Tocris Bioscience) and

refreshed on days 6–9. iECs were isolated on day 10 using MACS sorting using MiniMACS separator and kit (Miltenyi Biotec). Sorting was performed using CD34 MicroBead kit (Miltenyi Biotec).
iECs from cryo-preserved batches were used in all further experiments.
hiPSC-EC cells were thawed, resuspended in complete Endothelial cell serum-free medium (Gibco), and plated on a 0.1% gelatine-coated culture flask, as previously described. Cells were used for experiments when nearly confluent by visual inspection, typically on day 4. Cells were harvested using TrypLE™ according to the manufacturer's instructions. All iECs were used at passage #2.

*CRISPR-mediated gene editing.*
To generate the isogenic line for *COL4A1*$^{G755R}$ (iCOL4A1) and *COL4A2*$^{G702D}$ (iCOL4A2), a CRISPR-gene editing method was performed using single guide synthetic RNA (sgRNA; Synthego), SpCas9 protein (Biochemistry Department, University of Cambridge), and a 90-nt single-stranded oligodeoxynucleotide (ssODN; IDT) for homology-directed repair (**Table S2**). To avoid ssODN cleavage by Cas9, a silent mutation was introduced in the NGG codon upstream of the correction site (**Figure S1A, B**). For gene targeting, 200,000 cells were electroporated with Cas9/sgRNA together with ssODN using the Amaxa 4DNucleofector CA-137 program code (Lonza). Transfected cells were plated onto vitronectin coated-plates in TeSR™-E8 media with 10 μM Y-27632 and CloneR (STEMCELL Technologies). After 48h, the pool of transfected cells was sequenced to test recombination efficiency. Positive clones were selected by serial dilution and manual selection. Two sub-clones for each isogenic line were used for this study.

*Quantitative real-time polymerase chain reaction.*
Complementary DNA (cDNA) was synthesized from 250 ng total RNA using the Maxima First Strand cDNA Synthesis Kit (Thermo Fisher Scientific). Quantitative realtime polymerase chain reaction (qRT-PCR) mixtures were prepared with the FAST-SYBR Green Master Mix (Thermo Fisher Scientific) and analysed using the QuantStudio 7 Flex (Applied Biosystems, Thermo Fisher). CT values were normalised to housekeeping genes, GAPDH and PDGB using standard curve system. Primer sequences are listed in **Table S3.**

*Immunofluorescence staining and quantification*
Adherent cells were fixed using 4% PFA (Boster) for 5 minutes at RT (hiPSC and MC) or 100% ice cold Methanol (BMEC and iECs) for 15 minutes at -20°C and then washed 3 times with 1X PBS containing Calcium and Magnesium (Oxoid). Cells were permeabilised with 0.05% Triton X-100 (Sigma) in PBS and blocked with PBS +3% BSA or 10% FBS for 60 min at RT. For the detection of cell surface collagen IV, MC were incubated in blocking solution without permeabilisation. Primary antibodies (1:200; **Table S4**) incubations were performed at 4°C overnight and Alexa Fluor tagged secondary antibodies (1:500, Molecular Probes Invitrogen) and DAPI (Sigma-Aldrich) applied for 1 hour at room temperature the following day. Images were acquired on a Zeiss LSM 700 confocal and Leica TCS SP5 microscopes and analysed with Fiji-ImageJ software.
Quantification of fluorescence intensity for collagen IV was performed by taking the mean pixel intensity (Integrated Density, threshold 75-170) relative to the number of DAPI-positive cells, from an average of 3-5 fields of view from the same well. For tight junctions quantification in hiPSC-BMEC and iECs, following immunostaining

with occludin or claudin-5 antibodies, cells that lacked at least one continuous junction or show one frayed area were classified as discontinuous as previously described (Lee et al., 2018). Images were processed in Fiji-ImageJ software with a minimum of 5 fields with approximately 30 cells/field from three separate differentiations were quantified and all experimental groups remained blinded until completion of the study. All images are representative images.

*Western blotting*
Cells were lysed in RIPA buffer with added phosphatase inhibitor cocktail (Sigma) and protease inhibitor cocktail (Sigma) on ice for 15min. Protein content was quantified by Pierce Bicinchoninic Acid (BCA) Protein Assay Kit (Thermo Fisher Scientific). Samples (20ng) was resolved by electrophoresis on 10-15% Tris-HCl precast sodium dodecyl sulfate (SDS)-polyacrylamide gel (Bio-Rad), then transferred to polyvinylidine difluoride membranes (PVDF; Millipore). Membranes were blocked for 1 h at room temperature with 5% BSA in Tris-Buffered Saline containing 0.1% Tween-20 (TBS-T; Sigma) and incubated overnight with primary antibodies (**Table S4**) at 4°C. Membranes were washed with TBS-T, incubated with horseradish peroxidase (HRP)-conjugated secondary antibodies for 1 h at room temperature and developed with the Pierce ECL2 western blotting substrate (Thermo Fisher Scientific) using Gel Doc™ XR+ system (BioRad). The ImageLab™ Software (v5.2, BioRad) High Resolution programme with Signal Accumulation Mode was used to capture images at incremental exposure times. Anti-β-actin and GAPDH antibody was used as control for equal loading and transfer of the samples.

*BMEC/iECs Flow Cytometry*
Confluent wells were disassociated using Accutase and filtered to a single cell solution through a 40μm cell strainer (Corning®, Fisher Scientific). The cell suspension was fixed using Fixation/Permeabilisation solution (BD Biosciences) at 4ºC for 10 minutes and washed twice in PBS + 10% FBS. Cells were re-suspended with primary antibodies or pre-conjugated antibodies and incubated for 30 minutes at 4ºC. Cells were then resuspended in PBS and measured with a BD LRSFortessa or Canto II Flow Cytometer (BD Bioscience). Flow cytometric data were analysed with FCSalyzer 0.9.15-alpha software.

*Scratch migration assay*
Cells were plated onto 12-well plates and allowed to form a confluent monolayer. The cell monolayer was then scratched in a straight line to make a ‘‘scratch wound’’ with a 1-mL pipette tip. Cells were maintained in DMEM and images of the closure of the scratch were captured at different time points as indicated. Cells were tracked using the Wound_healing_size_tool macro for Fiji/ImageJ.

*Mouse aorta dissection and analysis*
Animal studies were performed in accordance with UK Home Office regulations (Project license 70/8604). Animals were sacrificed using an increasing gradient of $CO_2$ according to UK Home Office guidelines, and the thoracic aorta was collected and snap frozen on dry ice. Tissue samples were homogenized using steel beads (Qiagen) in TissueLyser (Qiagen) in RIPA buffer containing protease (Complete Mini, Roche) and phosphatase inhibitors (PhosSTOP, Roche). Protein concentrations were assessed via Pierce BCA Protein Assay (ThermoFisher) and protein were separated by SDS-PAGE (Mini-Protein Biorad). Membranes were blocked with 5% milk before incubation with primary and secondary antibodies and development

using chemiluminescense (Millipore). Protein levels were corrected for Coomassie staining of total protein gels ran or protein stain on membrane (Memcode, Pierce). Densitometry was performed using Image J.

### Table S1. iPSC lines

| Name | Vendor or Source | Sex and age | Individual | URL and Reference | Reprogram method/gene editing |
|---|---|---|---|---|---|
| WT1 (HPSI0414i-seru_7) | HIPSCI Consortium | F 65-69 | | https://www.hipsci.org/lines/#/lines/HPSI0414i-seru_7 | Sendai virus |
| WT2 (HPSI0314i-sojd_3) | HIPSCI Consortium | F 45-49 | | https://www.hipsci.org/lines/#/lines/HPSI0314i-sojd_3 | Sendai virus |
| WT3 (HPSI0214i-wibj_2) | HIPSCI Consortium | F 55-59 | | https://www.hipsci.org/lines/#/lines/HPSI0214i-wibj_2 | Sendai virus |
| *COL4A1*$^{G755R}$ Clone 4 and 5 | iPS Core Facility, Cambridge | F 65 | SVD patient | (Shah et al) | Sendai virus |
| *COL4A2*$^{G702D}$ | - | M 75 | Father of SVD patient | (Murray et al) | Sendai virus |
| iCOL4A1 clone 6 and 11 | - | F 65 | | - | CRISPR/Cas9 edited |
| iCOL4A2 clone 14 and 17 | - | M 75 | | - | CRISPR/Cas9 edited |

### Table S2. CRISPR sgRNA guide and ssODN

| **Gene/Mutation** | **Sequencing primers sequence 5'-3'** | **gRNA** | **Donor sequence 5'-3' (ssODN)** **Corrected base** **Mutated PAM** |
|---|---|---|---|
| *COL4A1*$^{G755R}$ | GCTTGAAAAGGGTTGAGCAG | CCGGCATTCCTGGCACACCC | G*A*C*TCAAAGGTTTGCCAGGTCTTCCCGGCATTCCTGGCACACCCGGAGAGAAGGGGAGCATTGGGGTACCAGGCGTTCCTGGAGAAC*A*T*G |
| *COL4A2*$^{G702D}$ | TCCAGTCCGTAAACAGGATTT | CGAAGCCUGGGAUUCCUCGG | G*C*C*TGATGTGGTTTGTGGTTTATTTGGTTATTTAGGTGCCAAAGGTCTCCG |

| | | | |
|---|---|---|---|
| | | | AGGAATCCCAGGCTTCG CAGGAGCTGATGGAGGA C*C*A*G |

### Table S3. qRT-PCR Primers sequences

| Gene Target | Forward Sequence 5’-3’ | Reverse Sequence 5’-3’ |
|---|---|---|
| *GAPDH* | AACAGCCTCAAGATCATCAGC | GGATGATGTTCTGGAGAGCC |
| *HMBS* (PBGD) | GGAGCCATGTCTGGTAACGG | CCACGCGAATCACTCTCATCT |
| *POU5F1* (OCT4) | AGGGCAAGCGATCAAGCA | GGAAAGGGACCGAGGAGTA |
| *SOX2* | ATGCACCGCTACGACGTGA | CTTTTGCACCCCTCCCATTT |
| *NANOG* | ACTAACATGAGTGTGGATCC | TCATCTTCACACGTCTTTCAG |
| *PECAM1* | CAGGCGCCGGGAGAAGTGAC | CGTCCAGTCCGGCAGGCTCT |
| *CD34* | CACAGGAGAAAGGCTGGGCGA | TGGCCGTTTCTGGAGGTGGC |
| *OCLN* | GGAGTGAACCCAACTGCTCA | CTCCTGGGGATCCACAACAC |
| *CDH5* | GGTCAAACTGCCCATACTTG | CGCAATAGACAAGGACATAACAC |
| *CLDN5* | CAGTACCGCAGGAAGAGGAG | ATCCCATGGCAAACAGAGAG |
| *CHD5* | CTCTGGGAGTGAGTGGAAGC | CCTGAGGATGATGGGAAAGA |
| *MMP2* | TCTCCTGACATTGACCTTGGC | CAAGGTGCTGGCTGAGTAGATC |
| *MMP9* | TTGACAGCGACAAGAAGTGG | GCCATTCACGTCGTCCTTAT |
| *MMP14* | CAGAGAAGGCACACAAACGA | CACTGGTGAGACAGGCTTGA |
| *CNN1* | GTCCACCCTCCTGGCTTT | AAACTTGTTGGTGCCCATCT |
| *P75* | ACAAGACCTCATAGCCAGCAC | CTGTTGGCTCCTTGCTTGTTC |
| *CSPG4* (NG2) | TTCCAGCTGAGCATGTCTGA | TCCTCCCGATCTGAAACCAC |
| *PDGFRB* | GCTTAAATCCACAGCCCGCA | AGGTAGTCCACCAGGTCTC |

### Table S4. Primary antibodies

| Target antigen | Species | Supplier | Catalogue number | Use |
|---|---|---|---|---|
| OCT3/4 | Mouse | Santa Cruz | SC-5279 | Immunofluorescence |
| SOX2 | Mouse | Abcam | sc-21705 | Immunofluorescence |
| TRA-1-60 | Rabbit | R&D Systems | AF2018-SP | Immunofluorescence |
| GATA-4 | Mouse | Santa Cruz | sc-25310 | Immunofluorescence |
| Brachyury | Mouse | Santa Cruz | sc-166962 | Immunofluorescence |
| occludin | Mouse | Thermo Fisher | 331500 | Immunofluorescence Western blotting |
| claudin-5 | Rabbit | Abcam | ab15106 | Immunofluorescence |

| | | | | |
|---|---|---|---|---|
| | | | | Western blotting |
| P75 | Rabbit | Abcam | ab8874 | Immunofluorescence |
| Smooth Muscle Actin | Mouse | Agilent | M085101-2 | Immunofluorescence |
| SM22 | Rabbit | Abcam | ab14106 | Immunofluorescence |
| Calponin | Mouse | Sigma-Aldrich | C-2687 | Immunofluorescence |
| NG2 | Rabbit | Sigma-Aldrich | AB5320 | Immunofluorescence |
| Collagen IV | Rabbit | Abcam | ab6586 | Immunofluorescence |
| claudin-5 pre-conjugated AF488 | Mouse | Thermo Fisher | 352588 | Flow cytometry |
| CD144 (VE-cadherin) APC-conjugated | Mouse | Thermo Fisher | 17-1441-80 | Flow cytometry |
| IgG1 Isotype Control FITC-conjugated | Mouse | Thermo Fisher | GM4992 | Flow cytometry |
| IgG1kappa Isotype Control APC-conjugated | Mouse | R&D Systems | IC002A | Flow cytometry |
| Annexin V-488 | - | Life technologies | V13241 | Flow cytometry |
| Propidium Iodide (PI) | - | Life technologies | | Flow cytometry |
| β-Actin | Mouse | Sigma-Aldrich | A1978 | Western blotting |
| MMP14 | Rabbit | Abcam | ab51074 | Western blotting |
| GAPDH | Mouse | Abcam | Ab8245 | Western blotting |

### Table S5. List of identified ECM differentially expressed genes (DEGs) in COL4A1/A2 vs isogenic MC.

| GENE | log2FoldChange | pvalue | padj | ECM |
|---|---|---|---|---|
| LAMA3 | 3.161270494 | 1.55E-09 | 6.08E-06 | 1 |
| ANK2 | 0.882039274 | 3.05E-08 | 4.18E-05 | 1 |
| ANKS1B | 3.778361044 | 8.95E-08 | 0.00010012 | 1 |
| NCAM1 | 4.111743201 | 1.83E-07 | 0.000151178 | 1 |
| DLGAP1 | 3.873590087 | 3.97E-07 | 0.000248996 | 1 |
| CORO2B | 1.159921465 | 4.78E-07 | 0.000277198 | 1 |
| ELMO1 | 3.045570878 | 1.93E-06 | 0.000718439 | 1 |
| MMP15 | 2.301754076 | 6.26E-06 | 0.001690658 | 1 |

| | | | | |
|---|---|---|---|---|
| **ZNF536** | 4.467677982 | 8.57E-06 | 0.002165223 | 1 |
| **USO1** | -1.52E-06 | 1.53E-05 | 0.003343641 | 1 |
| **ITPR2** | 0.456699641 | 1.54E-05 | 0.003343641 | 1 |
| **CNNM1** | 2.145289403 | 2.09E-05 | 0.004182211 | 1 |
| **AP3B2** | 1.912811981 | 4.83E-05 | 0.007132086 | 1 |
| **PRODH** | 3.358169282 | 5.14E-05 | 0.007253538 | 1 |
| **LONRF2** | 1.866100023 | 7.43E-05 | 0.008967966 | 1 |
| **SHROOM2** | 1.13143841 | 8.98E-05 | 0.010212208 | 1 |
| **ASAH1** | 0.643114655 | 9.48E-05 | 0.010539334 | 1 |
| **KCNA2** | 2.927873943 | 9.48E-05 | 0.010539334 | 1 |
| **GPR158** | 2.676727681 | 9.63E-05 | 0.010554064 | 1 |
| **CADM2** | 3.114526758 | 0.000128668 | 0.0130916 | 1 |
| **STX12** | 0.361590069 | 0.000145522 | 0.013722038 | 1 |
| **LIMCH1** | 2.611981232 | 0.000144025 | 0.013722038 | 1 |
| **TAGLN3** | 2.709206238 | 0.000143082 | 0.013722038 | 1 |
| **LGR5** | 3.343828338 | 0.000141258 | 0.013722038 | 1 |
| **LAMP5** | 1.56E-06 | 0.000150876 | 0.013940475 | 1 |
| **CNTN1** | 2.641453082 | 0.000151246 | 0.013940475 | 1 |
| **GAD1** | 1.715952087 | 0.000155125 | 0.014131677 | 1 |
| **MMP2** | -0.716063799 | 0.000158244 | 0.014324275 | 1 |
| **SH3GL2** | 1.813414845 | 0.000159181 | 0.014324275 | 1 |
| **ALDH1L1** | 2.529248317 | 0.000166382 | 0.014678375 | 1 |
| **PPP1R9A** | 1.487854112 | 0.000189881 | 0.016216903 | 1 |
| **NFS1** | 0.220249263 | 0.000192888 | 0.01633713 | 1 |
| **ABCB1** | 2.800720806 | 0.00024637 | 0.019301888 | 1 |
| **MMP7** | 3.976884934 | 0.000271229 | 0.020334379 | 1 |
| **FUBP1** | -0.948504724 | 0.00027813 | 0.020654138 | 1 |
| **MYOF** | 0.275804043 | 0.000310817 | 0.021645263 | 1 |
| **PPM1H** | 1.657361416 | 0.000309204 | 0.021645263 | 1 |
| **DOCK3** | 1.709308269 | 0.000309457 | 0.021645263 | 1 |
| **LAMA1** | 1.749551287 | 0.000398443 | 0.025722389 | 1 |
| **CLU** | 1.618955484 | 0.000405955 | 0.025962895 | 1 |
| **ATP6V1F** | 0.315755604 | 0.000411633 | 0.026112847 | 1 |
| **CASK** | 0.83790689 | 0.000478041 | 0.028434918 | 1 |
| **PECAM1** | -2.300979544 | 0.000540137 | 0.03033479 | 1 |
| **SELENBP1** | -2.96E-07 | 0.000657515 | 0.035701322 | 1 |
| **RTN1** | 2.234657952 | 0.000775382 | 0.039574786 | 1 |
| **DCLK1** | 1.305082133 | 0.000780584 | 0.039710944 | 1 |
| **ATP1A2** | 2.070141105 | 0.000849882 | 0.042029812 | 1 |
| **AAAS** | -0.234768613 | 0.000868548 | 0.042528978 | 1 |
| **RBFOX3** | 1.651824793 | 0.000894707 | 0.0433679 | 1 |
| **MMP24** | 1.725421609 | 0.000902287 | 0.0433679 | 1 |

| | | | | |
|---|---|---|---|---|
| **LGI3** | 1.232121367 | 0.000997875 | 0.046015926 | 1 |
| **COL4A6** | 1.377615567 | 0.001009755 | 0.046015926 | 1 |
| **SH3GL3** | 1.964934673 | 0.001032751 | 0.046234773 | 1 |
| **MMP9** | 1.964934673 | 0.001032751 | 0.046234773 | 1 |
| **SPTBN2** | 1.393768015 | 0.001080281 | 0.046790914 | 1 |
| **NBEA** | 0.708993913 | 0.001100329 | 0.047122048 | 1 |
| **SBSPON** | 1.624760968 | 0.001133237 | 0.0481538 | 1 |
| **MMP3** | 2.093771598 | 0.001175829 | 0.049394291 | 1 |
| **PRDX6** | 0.383723822 | 0.00119284 | 0.049694083 | 1 |

### Table S6. The Reactome pathways analysis of the identified ECM DEGs.

| Pathway identifier | Pathway name | pValue | Gene ID |
|---|---|---|---|
| **R-HSA-1592389** | Activation of Matrix Metalloproteinases | 1.7E-08 | MMP24;MMP7;MMP15;MMP2;MMP3;MMP9 |
| **R-HSA-1474244** | Extracellular matrix organization | 3.3E-08 | MMP24;MMP7;MMP15;LAMA1;MMP2;LAMA3;MMP3;COL4A6;PECAM1;CASK;NCAM1;MMP9 |
| **R-HSA-1474228** | Degradation of the extracellular matrix | 4.37E-07 | MMP24;MMP7;MMP15;MMP2;LAMA3;MMP3;COL4A6;MMP9 |
| **R-HSA-1442490** | Collagen degradation | 8.81E-07 | MMP7;MMP15;MMP2;MMP3;COL4A6;MMP9 |
| **R-HSA-2022090** | Assembly of collagen fibrils and other multimeric structures | 1.57E-05 | MMP7;LAMA3;MMP3;COL4A6;MMP9 |
| **R-HSA-373760** | L1CAM interactions | 3.17E-05 | LAMA1;CNTN1;NCAM1;ANK2;SH3GL2;SPTBN2 |
| **R-HSA-1474290** | Collagen formation | 1.24E-04 | MMP7;LAMA3;MMP3;COL4A6;MMP9 |
| **R-HSA-3000171** | Non-integrin membrane-ECM interactions | 1.91E-04 | LAMA1;LAMA3;COL4A6;CASK |
| **R-HSA-9022927** | MECP2 regulates transcription of genes involved in GABA signaling | 3.68E-04 | GAD1 |
| **R-HSA-3000157** | Laminin interactions | 4.16E-04 | LAMA1;LAMA3;COL4A6 |
| **R-HSA-6785807** | Interleukin-4 and Interleukin-13 signaling | 4.33E-04 | MMP7;MMP2;MMP3;MMP9 |
| **R-HSA-3000178** | ECM proteoglycans | 5.04E-04 | LAMA1;LAMA3;COL4A6;NCAM1 |
| **R-HSA-6806834** | Signaling by MET | 7.52E-04 | SH3GL3;LAMA1;LAMA3;SH3GL2 |
| **R-HSA-9009391** | Extra-nuclear estrogen signaling | 0.001760099 | MMP7;MMP2;MMP3;MMP9 |
| **R-HSA-70688** | Proline catabolism | 0.001955051 | PRODH |
| **R-HSA-9006934** | Signaling by Receptor Tyrosine Kinases | 0.002171418 | SH3GL3;DOCK3;LAMA1;LAMA3;ELMO1;ITPR2;MMP9;SH3GL2;ATP6V1F |
| **R-HSA-2214320** | Anchoring fibril formation | 0.002237685 | LAMA3;COL4A6 |
| **R-HSA-5578775** | Ion homeostasis | 0.003286605 | ITPR2;ATP1A2 |

| | | | |
|---|---|---|---|
| **R-HSA-8875360** | InlB-mediated entry of Listeria monocytogenes into host cell | 0.003548008 | SH3GL3;SH3GL2 |
| **R-HSA-422475** | Axon guidance | 0.005344994 | LAMA1;MMP2;CNTN1;NCAM1;ANK2;MMP9;SH3GL2;SPTBN2 |
| **R-HSA-6807004** | Negative regulation of MET activity | 0.006034786 | SH3GL3;SH3GL2 |
| **R-HSA-8876384** | Listeria monocytogenes entry into host cells | 0.006997575 | SH3GL3;SH3GL2 |
| **R-HSA-9675108** | Nervous system development | 0.007565737 | LAMA1;MMP2;CNTN1;NCAM1;ANK2;MMP9;SH3GL2;SPTBN2 |
| **R-HSA-8874081** | MET activates PTK2 signaling | 0.009685522 | LAMA1;LAMA3 |
| **R-HSA-445095** | Interaction between L1 and Ankyrins | 0.010270056 | ANK2;SPTBN2 |
| **R-HSA-182971** | EGFR downregulation | 0.012759693 | SH3GL3;SH3GL2 |
| **R-HSA-6807878** | COPI-mediated anterograde transport | 0.013357171 | USO1;ANK2;SPTBN2 |
| **R-HSA-9768919** | NPAS4 regulates expression of target genes | 0.015484716 | RBFOX3 |
| **R-HSA-8875878** | MET promotes cell motility | 0.01843598 | LAMA1;LAMA3 |
| **R-HSA-9609736** | Assembly and cell surface presentation of NMDA receptors | 0.021604588 | NBEA;CASK |
| **R-HSA-1500931** | Cell-Cell communication | 0.02404666 | CADM2;LAMA3;CASK |
| **R-HSA-1489509** | DAG and IP3 signaling | 0.024981882 | NBEA;ITPR2 |
| **R-HSA-3928665** | EPH-ephrin mediated repulsion of cells | 0.026746145 | MMP2;MMP9 |
| **R-HSA-888568** | GABA synthesis | 0.027188048 | GAD1 |
| **R-HSA-5576891** | Cardiac conduction | 0.03044768 | ITPR2;ATP1A2 |
| **R-HSA-8939211** | ESR-mediated signaling | 0.030796928 | MMP7;MMP2;MMP3;MMP9 |
| **R-HSA-6794361** | Neurexins and neuroligins | 0.031369098 | CASK;DLGAP1 |
| **R-HSA-9634815** | Transcriptional Regulation by NPAS4 | 0.031369098 | RBFOX3 |
| **R-HSA-9032759** | NTRK2 activates RAC1 | 0.031647996 | DOCK3 |
| **R-HSA-177929** | Signaling by EGFR | 0.032329074 | SH3GL3;SH3GL2 |
| **R-HSA-166665** | Terminal pathway of complement | 0.036087788 | CLU |
| **R-HSA-112043** | PLC beta mediated events | 0.038326293 | NBEA;ITPR2 |

| | | | |
|---|---|---|---|
| **R-HSA-199977** | ER to Golgi Anterograde Transport | 0.040039399 | USO1;ANK2;SPTBN2 |
| **R-HSA-2161517** | Abacavir transmembrane transport | 0.040507513 | ABCB1 |
| **R-HSA-164944** | Nef and signal transduction | 0.040507513 | ELMO1 |
| **R-HSA-447043** | Neurofascin interactions | 0.040507513 | CNTN1 |
| **R-HSA-166520** | Signaling by NTRKs | 0.041257591 | SH3GL3;DOCK3;SH3GL2 |
| **R-HSA-375165** | NCAM signaling for neurite out-growth | 0.041472104 | NCAM1;SPTBN2 |
| **R-HSA-936837** | Ion transport by P-type ATPases | 0.04362168 | ATP1A2 |
| **R-HSA-112040** | G-protein mediated events | 0.044711834 | NBEA;ITPR2 |
| **R-HSA-446107** | Type I hemidesmosome assembly | 0.049287121 | LAMA3 |
| **R-HSA-9032500** | Activated NTRK2 signals through FYN | 0.049287121 | DOCK3 |
| **R-HSA-216083** | Integrin cell surface interactions | 0.04976213 | COL4A6;PECAM1 |

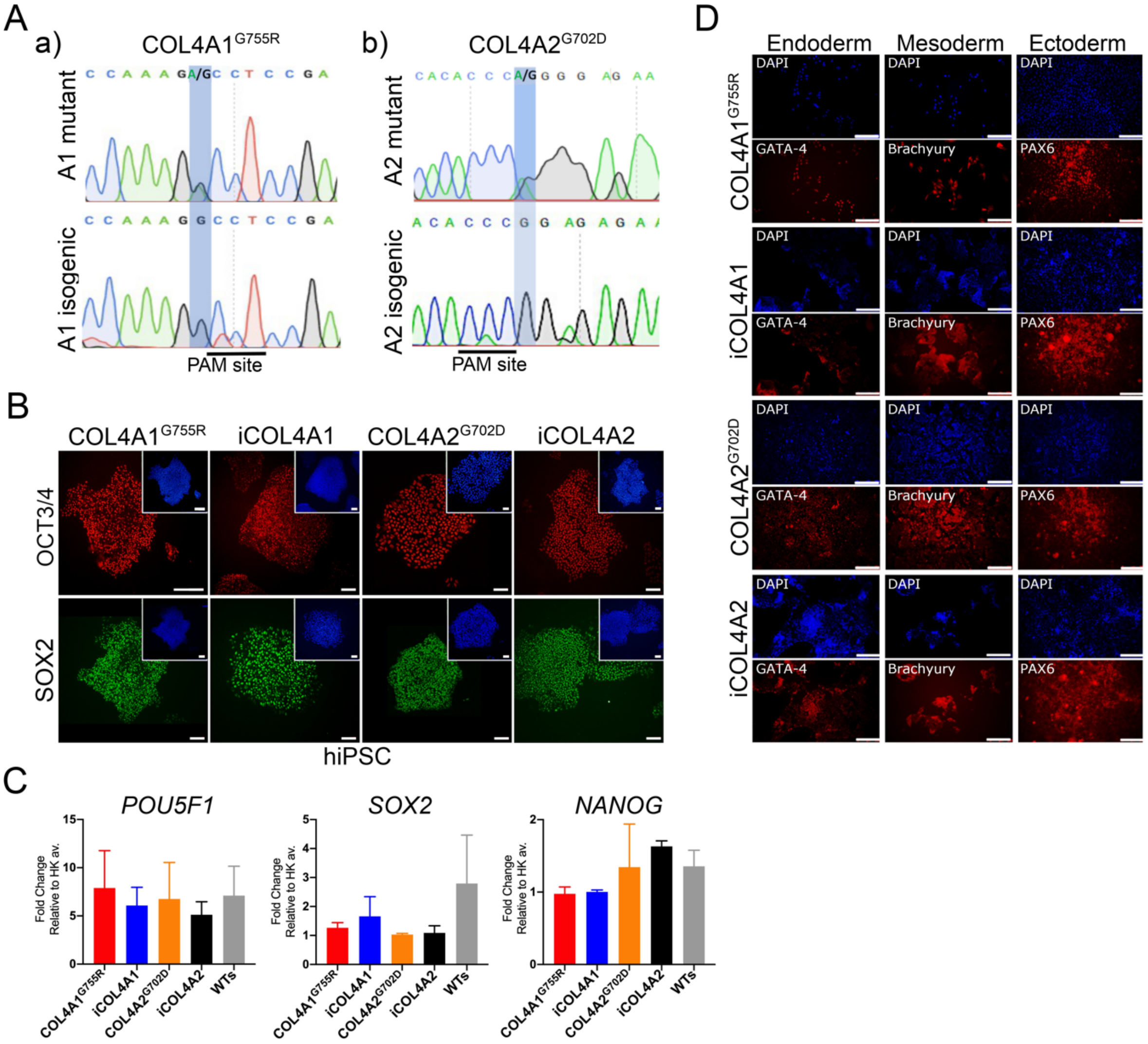


**Figure S1. hiPSC characterisation for COL4A1*G755R*, COL4A2*G702D* and isogenic lines.**

**A)** Sanger sequencing output for **(a)** *COL4A1* heterozygous mutation (G775R) and CRISPR-corrected isogenic A1 and **(b)** *COL4A2* heterozygous mutation (G702D) and CRISPR-corrected isogenic A2. **B)** Immunostaining analysis for hiPSC markers (OCT3/4 and SOX2) for *COL4A1G755R*, *COL4A2G702D* and isogenic A1 (iCOL4A1) and A2 (iCOL4A2); nuclei were stained with DAPI (insert); scale bar=100µm. **C)** RT-qPCR analysis for pluripotency markers expression (*POU5F1*, *SOX2* and *NANOG*); the results are presented as means ± SD of two independent sub-clones (three technical replicates). **D)** Immunostaining analysis for each of the three germ layers (GATA-4, endoderm; BRACHYURY, mesoderm; PAX6, ectoderm); nuclei were stained with DAPI; scale bar=100µm. hiPSC= induced pluripotent stem cells. PAM site= protospacer adjacent motif sequence.

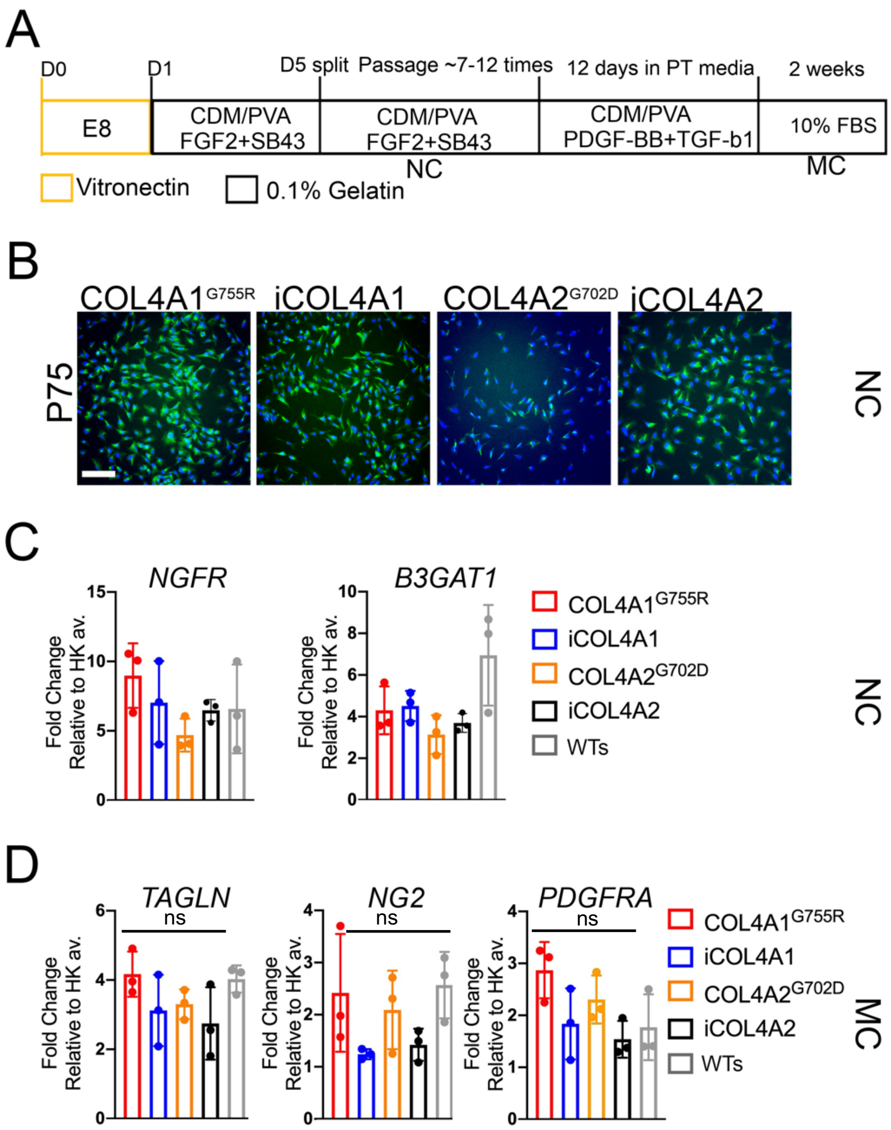


**Figure S2. Neural-crest derived smooth muscle cells differentiation and characterisation.**
**A)** Schematic of neural crest (NC) derived smooth muscle cells (MC) differentiation (Cheung *et al.)*. Characterisation of NC intermediate population by **(B)** immunostaining for specific marker (p75) and **(C)** RT-PCR for *NGFR* (P75) and *B3GAT1* (HNK1). **D)** RT-PCR for SMC markers (*TAGLN, NG2, PDGFRA*) for *COL4A1*$^{G755R}$, *COL4A2*$^{G702D}$, isogenic and WTs (three independent healthy control; **Table S1**) lines. NC=neural crest; MC=neural crest-derived smooth muscle cells. The results are presented as means ± SD of three independent experiments; ns, not significant. Statistical analysis was performed by 2-way ANOVA with Tukey's multiple comparison test.

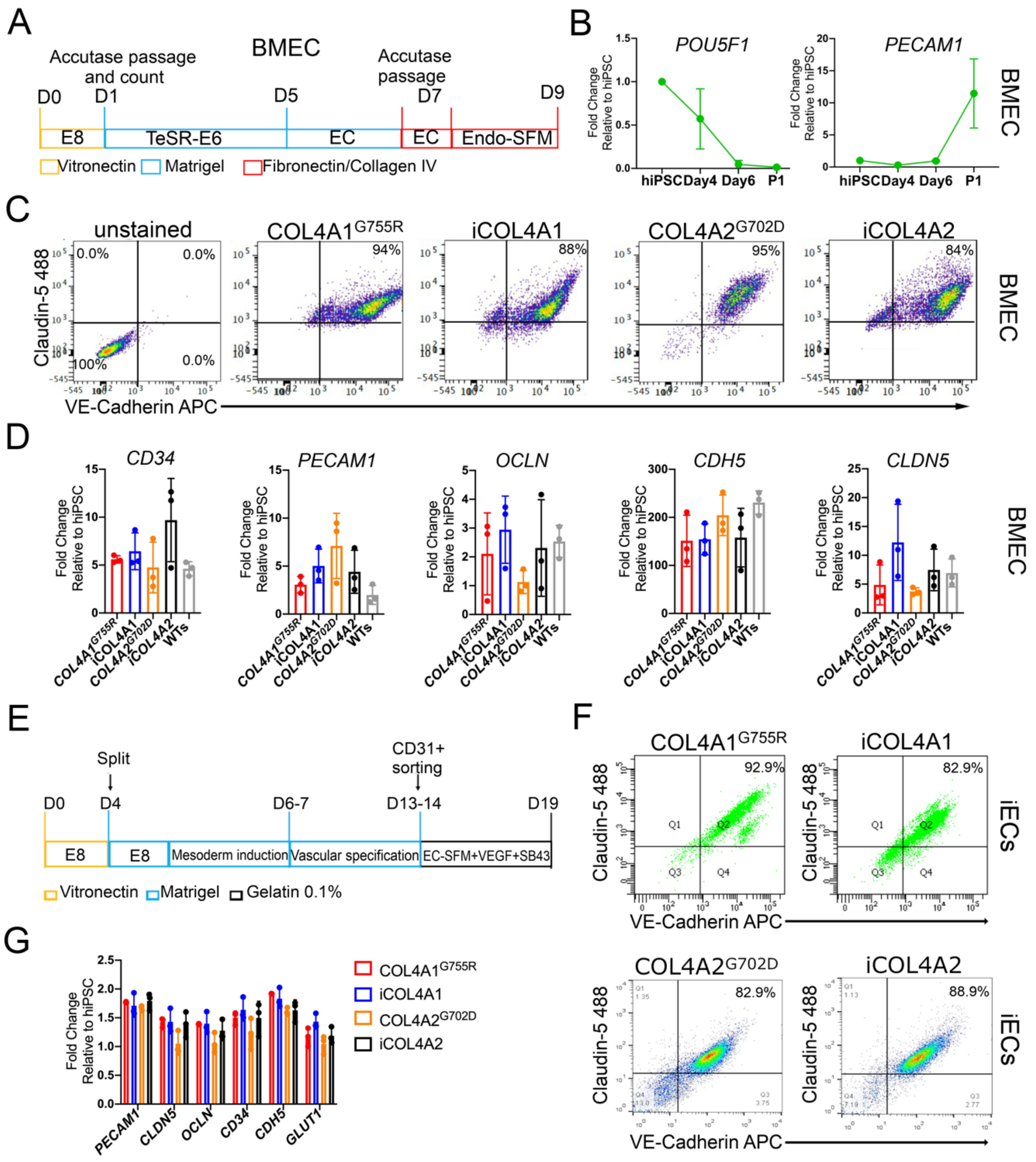

**Figure S3. hiPSC-derived brain microvascular endothelial-like cells (BMEC) and endothelial cells (iECs) differentiation and characterisation.**
**A)** Schematic of brain microvascular endothelial-like cells (BMEC) differentiation from hiPSC (Hollman *et al*). **B)** Representative time-course RT-qPCR of hiPSC-BMEC differentiation for pluripotent marker (*POU5F1*) and endothelial marker (*PECAM1*). **C)** Flow cytometric analysis of hiPSC-BMEC for VE-cadherin (APC conjugated) and claudin-5 (488 conjugated) for *COL4A1*$^{G755R}$, *COL4A2*$^{G702D}$ and isogenic lines. **D)** mRNA profile of hiPSC-BMEC by RT-qPCR for specific markers (*CD34, PECAM1, OCLN, CDH5 and CLDN5*). **E)** Schematic of hiPSC-endothelial cells (iECs; Orlova *et al*) differentiation and characterisation by **(F)** flow cytometry for VE-cadherin and claudin-5 double staining and (**G)** RT-qPCR for endothelial markes: *PECAM1, CLDN5, OCLN, CD34, CDH5 and GLUT-1*). The results are presented as means ± SD of three independent experiments.

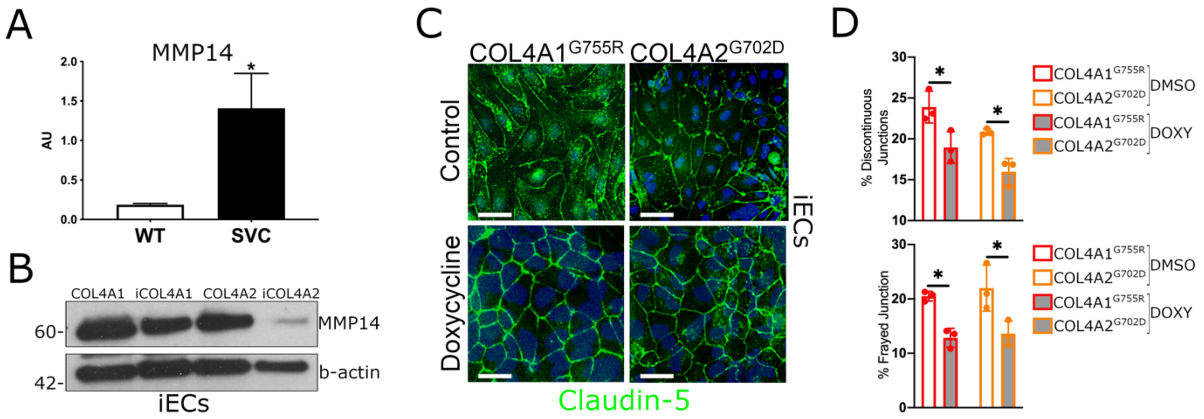


**Figure S4. MMP14 is increased in Col4a1 mouse aorta and hiPSC-derived ECs showing tight junction abnormalities.**

**A)** Total Mmp14 protein level found higher in Col4a1 mice aorta (n=6) compared to WT mice. **B)** Protein blot showing MMP14 upregulation in *COL4A1/A2* iECs compared to isogenics. **C-D)** Increased discontinuity and frayed junctions in *COL4A1/A2* hiPSC-ECs showed by staining for claudin-5 is reverted upon doxycycline treatment (DOXY). The results are presented as means ± SD of three independent experiments. $^{*}P < 0.05$. Statistical analysis was performed by 2-way ANOVA with Tukey's multiple comparison test.
